# Systemic AAV delivery of a calcium indicator in marmosets: functional validation in visual area MT

**DOI:** 10.64898/2026.01.23.701367

**Authors:** Penny-Shuyi Chen, Declan P Rowley, Mitchell Rudd, Allison Laudano, Andrew P Villa, Fernando Garcia, Hong-Wei Dong, Timothy F Shay, Alexander C Huk, Andrew D Steele, Joseph Wekselblatt

## Abstract

Functional optical imaging in nonhuman primates provides an important complement to electrophysiological approaches in neuroscience research, but its broader use has been limited by challenges in achieving large-scale, homogeneous expression of genetically encoded reporters, and imaging accessibility in species with gyrencephalic brains with sulci and fissures (e.g., rhesus macaques). Specifically, conventional local intracortical viral injections are invasive and often produce spatially restricted or heterogeneous expression, constraining population-level analyses. Here, we show that systemic intravenous delivery of an adeno-associated virus (AAV) capsid engineered for enhanced blood-brain barrier crossing, AAV.CAP-B10, supports robust and widespread expression of a calcium indicator CAaMP8s in the common marmoset. Intravenous delivery in two marmosets resulted in widespread cortical expression. Using a large cranial window over extrastriate visual area MT (and its satellite areas), we performed widefield single-photon imaging and two-photon cellular-resolution imaging in awake,behaving marmosets to functionally validate activity in this well studied primate visual-motion sensitive cortical area. Population level responses to visual motion and spatial organization measured with widefield imaging, as well as single-cell level motion direction tuning measured with two-photon imaging, were consistent with canonical properties of MT reported in previous electrophysiological studies. Quantitative analyses of lightsheet imaging after whole hemisphere brain clearing further confirmed the broad expression of GCaMP in both cortical and subcortical areas. Together, these results indicate that systemic delivery using AAV.CAP-B10 provides a minimally invasive approach for robust multi-scale functional optical imaging in awake, behaving marmosets.

## Introduction

Understanding the functional organization of the primate cortex requires methods capable of sampling neural activity across large spatial extents while preserving cellular resolution (Clough & Chen, 2019, 2019; Trautmann et al., 2023; Watakabe et al., 2023; Weisenburger & Vaziri, 2018). Primate visual area MT has served as a canonical model for studying cortical representations of motion, with decades of work characterizing its motion tuning properties, columnar organization, and population dynamics using electrophysiological techniques (Born & Bradley, 2005; Czuba et al., 2014; DeAngelis et al., 1998; Huk et al., 2002; Ilg, 2008; R. Maunsell, 1987; Rudolph & Pasternak, 1999; S. G. Solomon & Rosa, 2014; Z.-X. Xu & DeAngelis, 2025). While these approaches have provided detailed insight into single-neuron response properties, they inherently sample sparse neuronal populations and offer limited access to large-scale spatial organization (Moreaux et al., 2020; Schröter et al., 2025; Siegle et al., 2021; Steinmetz et al., 2018). Optical imaging approaches provide a complementary perspective, enabling simultaneous measurements across broad cortical regions and, at higher resolution, across large populations of individual neurons (Ebina et al., 2018; Mulholland et al., 2024; Nietz et al., 2022; Ohki et al., 2005; Sadakane, Masamizu, et al., 2015; Sadakane, Watakabe, et al., 2015; Song et al., 2024; Zhang et al., 2024).

In nonhuman primates, the application of functional imaging has been constrained by challenges associated with the delivery of genetically encoded activity indicators (Broussard et al., 2014; Campos et al., 2023; El-Shamayleh et al., 2016; L. Li & Liu, 2023; Park et al., 2016; Seidemann et al., 2016). Most studies rely on local intracortical viral injections, which are invasive, often require multiple penetrations to achieve sufficient coverage, and can produce heterogeneous expression patterns (Daci & Flotte, 2024; Fischell & Fishman, 2021; Lowery & Majewska, 2010; Zhou et al., 2022). These limitations are particularly restrictive for widefield imaging and longitudinal studies, where uniform and stable expression across large cortical regions is critical for interpreting population-level signals.

Recent advances in adeno-associated virus (AAV) engineering have produced capsid variants capable of crossing the blood–brain barrier (BBB) following systemic administration (Agbim et al., 2025; Carneiro & Schaffer, 2024; Challis et al., 2022; M. R. Chuapoco et al., 2023; Goertsen et al., 2022; Lopez-Gordo et al., 2024; Suarez-Amaran et al., 2025). Such vectors enable noninvasive, brain-wide delivery of genetic payloads, reducing the spatial and logistical constraints of local injections while improving the uniformity of reporter expression. In the common marmoset (*Callithrix jacchus*)— a lissencephalic New World primate increasingly used in systems neuroscience— systemic viral delivery is particularly attractive due to the accessibility of large cortical regions and the feasibility of chronic optical imaging (D’Souza et al., 2021; Ebina et al., 2018; Hung et al., 2015; Kishi et al., 2014; Mitchell et al., 2014; Sadakane, Masamizu, et al., 2015; S. G. Solomon & Rosa, 2014). However, while systemic delivery has been shown to produce widespread expression, relatively few studies have provided functional validation of genetically encoded calcium indicators delivered using this approach in non-human primates (M. Chuapoco et al., 2023, 2023).

In the primate visual system, area MT serves as a major cortical locus for the processing of motion signals that support perception, eye movements, and visually guided behavior (Born & Bradley, 2005; DeAngelis et al., 1998; Ilg, 2008; Lisberger & Movshon, 1999; J. H. Maunsell & Van Essen, 1983; J. Maunsell & Van Essen, 1983; Moutoussis & Zeki, 2008; S. G. Solomon & Rosa, 2014). In both macaques and humans, MT and its associated network (the “MT complex”) are strongly linked to motion perception, motion-induced eye movements, and perceptual decisions based on motion and depth (Bradley et al., 1998; K. Britten et al., 1992; Heeger et al., 1999; Lisberger & Movshon, 1999; Moutoussis & Zeki, 2008; W. T. Newsome et al., 1989; Orban et al., 2003; Schiller & Lee, 1994). Lesions or dysfunction of MT in primates produce selective impairments in motion perception, underscoring its functional specialization (Groh et al., 1997; Murasugi et al., 1993; T. Newsome et al., n.d.; W. Newsome & Pare, 1988; Nichols & Newsome, 2002; Salzman et al., 1992). Overall, the MT complex, as a visual-motion processing hub that represents a highly specialized cortical region conserved across primate species, exhibits important functional specialization and hierarchical computations with behavioral relevance, and which lacks a clearly established homolog across other mammalian orders (Allman & Kaas, 1971; Huk et al., 2002; Huk & Heeger, 2002; Kaskan, 2010; Kaskan & Kaas, 2007, 2007; Lui & Rosa, 2015; Sereno et al., 2015).

Despite decades of electrophysiological study in macaque MT, important aspects of its functional organization remain difficult to assess using sparse sampling approaches. Electrode recordings provide detailed information about single-neuron response properties but offer limited access to the spatial organization of motion tuning across large cortical regions, including the layout of motion direction, speed and disparity columns (Born & Bradley, 2005; Harris et al., 2016; Siegle et al., 2021; Wei et al., 2020). Optical imaging approaches can, in principle, resolve these structures directly. However, their application in macaques, in addition to the difficulty of achieving large-scale, homogeneous expression of fluorescent indicators, has been also constrained by the optical imaging accessibility in gyrencephalic brains. Notably, macaque MT is embedded in the dorsal bank of the posterior superior temporal sulcus (STS) (Desimone & Ungerleider, 1986; Erickson et al., 1989). The sulcal boundaries and structure can make it hard to obtain continuous imaging field of view and therefore fragmentize functional maps, limiting the interpretation of cortical functional organization (Song et al., 2022a).

The common marmoset has emerged as a particularly well-suited primate model for bridging the gap between rodent optical imaging and human-relevant visual systems (Mitchell & Leopold, 2015; Sadakane, Masamizu, et al., 2015; Schiessl & McLoughlin, 2003; S. G. Solomon & Rosa, 2014; Song et al., 2022b, 2024; Yamamori, 2021). As a diurnal primate with a foveated, cone-dominated retina and a visual cortical hierarchy that includes a clearly defined MT complex, marmosets share key organizational and functional features of motion processing with macaques and humans (Chaplin et al., 2017; Cloherty et al., n.d.; Q. Li et al., 2025; Lui & Rosa, 2015; Pattadkal et al., 2023; S. S. Solomon et al., 2011). At the same time, the marmoset’s lissencephalic and relatively small cortex provides substantially improved optical access compared to gyrencephalic primates, enabling widefield and cellular-resolution imaging across large contiguous cortical regions. These anatomical features make marmosets uniquely compatible with advanced optical imaging approaches that are difficult to deploy in macaques, while preserving the primate-specific circuitry and visual computations that are absent or substantially transformed in rodents (Kell et al., 2023; Magrou et al., 2024; Mitchell & Leopold, 2015; Skibbe et al., 2023; S. G. Solomon & Rosa, 2014). In combination with recent advances in systemic viral delivery of genetically encoded calcium indicators, the marmoset model provides an opportunity to perform multi-scale functional imaging of primate visual cortex with spatial and temporal resolution beyond what is achievable with intrinsic imaging or sparse electrophysiological sampling.

Our motivation for this study is to bridge the gap between well-established physiological benchmarks derived from technically demanding electrophysiological measurements and the advanced tools needed to directly and comprehensively visualize them. Therefore, in this report, we asked whether systemic intravenous delivery of a BBB–crossing AAV could support functional calcium imaging in the marmoset visual cortex at both population and cellular scales. We packaged the calcium indicator GCaMP8s into AAV.CAP-B10 (hereafter referred to as CAP-B10), which crosses the BBB with high efficiency and has a strong tropism to neurons after intravenous administration (Goertsen et al., 2022). We administered CAP-B10-CAG-GCaMP8s intravenously in two common marmosets and we performed widefield single-photon imaging to identify a region with robust visual-motion responses as the putative MT complex through a large cranial window over the extrastriate visual cortex.. Two-photon imaging, sampling different sub-regions within the area identified by widefield, was conducted to assess motion direction tuning at cellular level. Together, these widefield and cellular measurements in the marmoset MT complex provide functional validation that systemically delivered GCaMP yields reliable functional signals consistent with established properties of MT proposed by previous electrophysiological studies. Here, we show that this proof of concept establishes a foundation for leveraging additional strengths of imaging approaches, including cell-type-specific measurements and spatially precise optical stimulation.

## Methods

### General approach

We packaged a genetically encoded calcium indicator (GCaMP8s) into an adeno-associated viral capsid engineered to (i) cross the blood–brain barrier, (ii) preferentially transduce neurons, and (iii) reduce off-target expression in peripheral organs, such as the liver (Goertsen et al., 2022). This construct (AAV.CAP-B10-CAG-GCaMP8s) was delivered via intravenous tail-vein injection to two common marmosets (*C. jacchus*).

Following systemic delivery and an appropriate expression period, we performed widefield single-photon (1P) and two-photon (2P) calcium imaging experiments in awake, behaving animals. Imaging was conducted through large cranial windows positioned over posterior extrastriate cortex, encompassing the expected location of visual area MT and nearby motion-sensitive regions. Full-field moving dot stimuli alternating between motion and stationary epochs were first used during widefield imaging to identify visual-motion–responsive regions of interest (ROIs). The ROI exhibiting robust, stimulus-locked responses to motion was identified as the MT complex.

Additional motion stimuli were then used to characterize population-level functional properties of the MT complex, including direction tuning, speed tuning, and retinotopy, using widefield imaging. Two-photon imaging was performed in separate sessions using the same functional mapping stimuli, with cellular-resolution recordings obtained from multiple smaller ROIs within the MT complex identified by widefield imaging.

After completion of in vivo imaging experiments, brain tissue was processed for whole-brain clearing and imaged with light-sheet microscopy to independently assess the spatial extent and uniformity of GCaMP expression. In this methodology-oriented report, we focus on functional validation of systemically delivered GCaMP in awake marmoset MT by demonstrating robust population-level visual-motion responses with widefield imaging, direction-selective responses at the single-neuron level with two-photon imaging, and widespread cortical expression confirmed ex vivo with light-sheet imaging. Definitive identification of MT from the greater MT complex, as well as detailed quantitative analyses of MT’s functional organization (such as comprehensive assessments of direction tuning, speed tuning, and retinotopic organization), will be presented in a subsequent report.

### Animals

Two adult common marmosets (one male–age= 2 years and 1 month old, and one female–age=2 years and 4 months old by the day of imaging chamber surgery) were used in this study for the functional imaging experiments. One adult marmoset (male–age= 3 years and 2 months old, at time of IV injection with 3 months of expression allowed before histology) was used for the administration of CAP-B10-CAG-smV5-GFP virus. Animals were housed and cared for in accordance with NIH and USDA guidelines, and all procedures were approved by the UCLA Institutional Animal Care and Use Committee.

### Viral construct and production

We used a recombinant AAV engineered to cross the BBB and preferentially transduce neurons in the marmoset central nervous system (Goertsen et al., 2022). The calcium indicator GCaMP8s was expressed under the control of the CAG promoter and also included a Woodchuck Hepatitis Virus Posttranscriptional Regulatory Element (WPRE) to enhance expression. This construct was triple transfected into HEK cells along with pHelper and capsid plasmid encoding CAP-B10 and other elements needed for AAV packaging following established protocols (Challis et al., 2019). Vector genome copies were titered using digital PCR (AbsoluteQ system with Taqman ITR probes purchased from Thermofisher).

### Intravenous viral delivery

Animals were first sedated with Alfaxan (5-10 mg/kg) and midazolam (0.25 mg/kg) delivered intramuscularly via the same syringe. Then the virus was delivered slowly via tail-vein injection over approximately 10 minutes. Each animal received approximately 2.8 × 10¹³ vector genomes in a total volume of approximately 1.0 mL. Animals also received 4 days of oral Dexamethasone (0.25 mg/kg) to reduce possible autoimmune responses. Animals recovered without complication and were returned to their home cages after the procedure.

### Cranial imaging window and head restrainment implantation

For both animals, a large cranial imaging window, providing optical access to a circular region approximately 7mm in diameter, was implanted over the right hemisphere’s MT complex, based on the expected anatomical location of area MT, with the window planned to be centered in stereotaxic coordinates at Animal 1 = (+2.5mm AP interaural, +10.2mm ML), Animal 2 = (+1.14mm AP interaural, +10.24mm ML) (Liu et al., 2018) (see Figure 1C for Animal1’s MT window). The exact location of the imaging window center was adjusted during surgery to best accommodate the skull curvature for window chamber placement. The window chamber was adapted and modified from previously described designs developed for chronic imaging in marmoset (Pattadkal & Priebe, 2025). After a craniotomy was made, a titanium chamber was secured to the skull. Metabond and dental acrylic were applied to further secure the chamber placement. Then, a durotomy was carefully performed, followed by the placement of the chamber, which holds a transparent glass window to allow for functional imaging. After the imaging window was implanted, head restraint including a halo-style headplate (notched to allow for the placement of the chamber on the right hemisphere) and a headpost were implanted to ensure imaging stability during the experiments. Animals recovered without complications after the implantation surgery, and returned to their home cage. The animals were monitored for 10 days, and medication was administered in accordance to the research protocol and daily assessment (Meloxicam 0.1mg/kg; Dexamethasone 1.5mg/kg; Famotidine 0.8mg/kg; Cerenia 1mg/kg; all meds were given PO, sid). Imaging window clarity and GCaMP expression quality were assessed before functional mapping experiments began.

**Figure 1.**
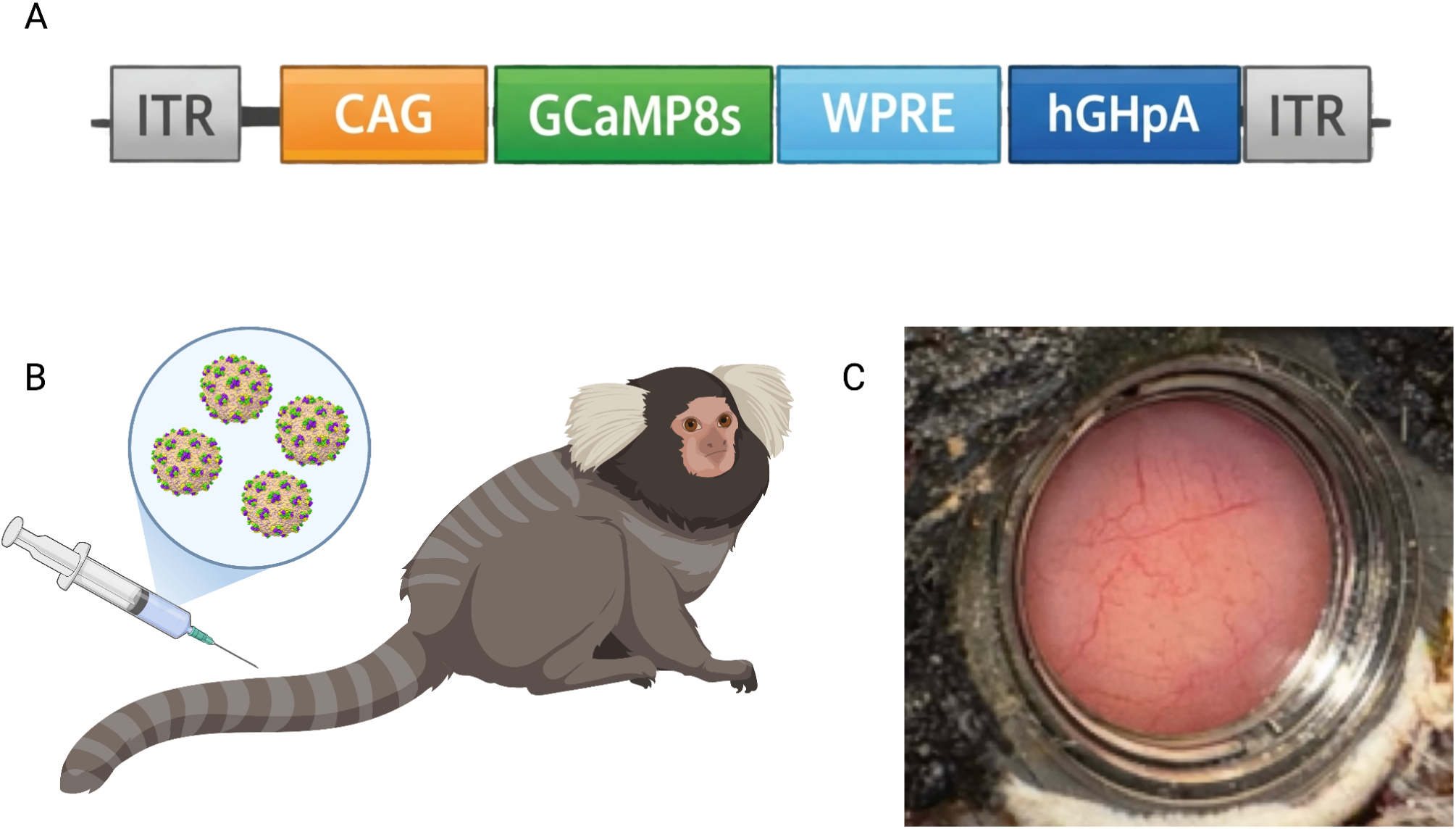
Systemic AAV delivery enables large-scale optical access to marmoset cortex. (A) Schematic of the viral construct used in this study (AAV9-CAP-B10-CAG-GCaMP8s). (B) Schematic of intravenous delivery of the BBB-crossing capsid via tail-vein injection. (C) Cranial window implanted posterior to the lateral/temporal sulcus and approximately centered on the expected location of area MT in marmoset.

### Imaging experiment rig instrumentation

The optical imaging experiment rig is shown in Supplemental Figure 1 with labels to each part of the apparatus.

### Widefield one-photon imaging

Widefield one-photon imaging was performed throughout the cranial window, and the acquisition mainly followed the protocol outlined in Couto et al. (2021), using a PCO panda 4.2 CMOS camera (Excelitas) and open-source software Labcams (Github link: https://github.com/jcouto/labcams). Frames (10Hz) were synchronized using a fast microcontroller (Teensy 4.1) with two LEDs illuminators alternately providing 470nm light for GCaMP excitation and 530nm light for hemodynamic correction. 530nm (THORLABS M530L4) was chosen as an alternative isosbestic point for equal absorption of oxygenated/deoxygenated hemoglobin (tracking blood volume) rather than violet as in Couto et al (2021) in order to minimize the cortex’s exposure to high energy light. Therefore, the functional images were acquired at 5Hz. As the 530nm light source could not pass through the same (498nm) dichroic filter as the blue excitation light (469nm bandpass filtered THORLABS MCWHL6 white LED), it was applied externally from the side rather than through the objective. The light was positioned to minimize the differences in illumination across the window due to its incident angle and the relative power (typically 10-20mW) was adjusted to approximately match the histograms between the two channels. A 2x/0.1NA objective (THORLABS) and a 4x/0.2NA objective (Nikon) were used for widefield imaging experiments.

### Two-photon imaging

Two-photon imaging was performed using a resonant scanning two photon microscope optimized for *in-vivo* optical imaging (Neurolabware, Los Angeles, CA) coupled to a Chameleon Vision II laser, tuned to 920nm, with a 16x/0.8NA objective (Nikon). Scanbox software (Dario Ringach / Neurolabware) in Matlab (Mathworks) was used for data acquisition. Images were acquired at 31.25 fps with bidirectional scanning and 796×796 pixels over a ∼800×800um field of view, using 65-250mW illumination power as measured at the front aperture of the objective. All recordings were targeted to layer 2/3 of MT (depth 150 – 350 microns).

### Optical imaging experiment platform for awake and behaving marmosets

For all imaging sessions, the animals were stably head-restrained by the halo-headplate holder and chaired in a cradle mounted to the body stabilizing system that damped the majority of up-down movements and translated any residual into x-y plane movements. This minimized the movement in the z-plane (up and down) during imaging.

### Visual display

All visual stimuli were presented on a linearized and calibrated flat fronto-parallel display (ASUS 27”), running at 240Hz with a viewing distance of 48cm and resolution at 2560×1440 pixels.

### Acclimation and Eye-tracking training

Before conducting imaging sessions, the animals first needed to get accustomed to being chaired at the imaging rig awake and then get trained for eye-tracking calibration. During these sessions, animals were head-fixed and trained to fixate and tested for visual acuity. Each session was controlled to be under 90 minutes. As the animal was fully trained, at the beginning of each imaging experiment session, the animal’s eye-tracking was re-calibrated and refined based on last calibration. High-resolution dual-purkinje image (DPI) binocular eye-tracking was applied to precisely register the eye positions and the eye-tracking data were acquired through OpenIris software at 500 Hz (Ressmeyer et al., 2025). Eye position was used to ensure accurate stimulus alignment, which is crucial for analyzing MT motion tuning properties (Bair & O’Keefe, 1998; Lisberger & Movshon, 1999; Price et al., 2005).

### MT complex functional mapping experiments

We conducted functional widefield one-photon and two-photon imaging (in separate sessions), with a battery of full-field moving dot stimuli, to assess the putative MT complex’s motion direction tuning, speed tuning and retinotopy. In this report’s method section, we focus on the stimuli used to 1) identify visual-motion sensitive areas within the cranial window (1P only), and 2) assess motion direction tuning (1P and 2P), with an emphasis on functionally validating the expression. The comprehensive description of the stimulus battery will be included in a separate report (in preparation) specifically investigating the visual-motion tuning properties and spatial organization of the functional structures across retinotopy in common marmoset area MT, at both population and cellular levels.

### Motion-stationary dot stimulus

In order to functionally select and assess visual-motion responsive ROIs within the cranial window, stimuli with alternating full-field dot motion (600ms) and stationary dots (600ms) periods were shown to animals (Figure 2A). For each frame, there were 18 large white dots, and each moved in one of the 16 motion directions (spaced in 22.5 deg steps across all 360 possible deg) at a randomly selected speed log spaced between 2 and 32 visual degrees per second. The lifetime of each dot was 50ms (12 frames at 240Hz). There were 20 motion-stationary cycles (1.2s x 20) in each trial, and 40 trials, with 2s long inter-trial-interval, were run for one session. This stimulus was used for widefield 1P imaging only, and the 2x/0.1NA lens was used to image the entire window.

**Figure 2.**
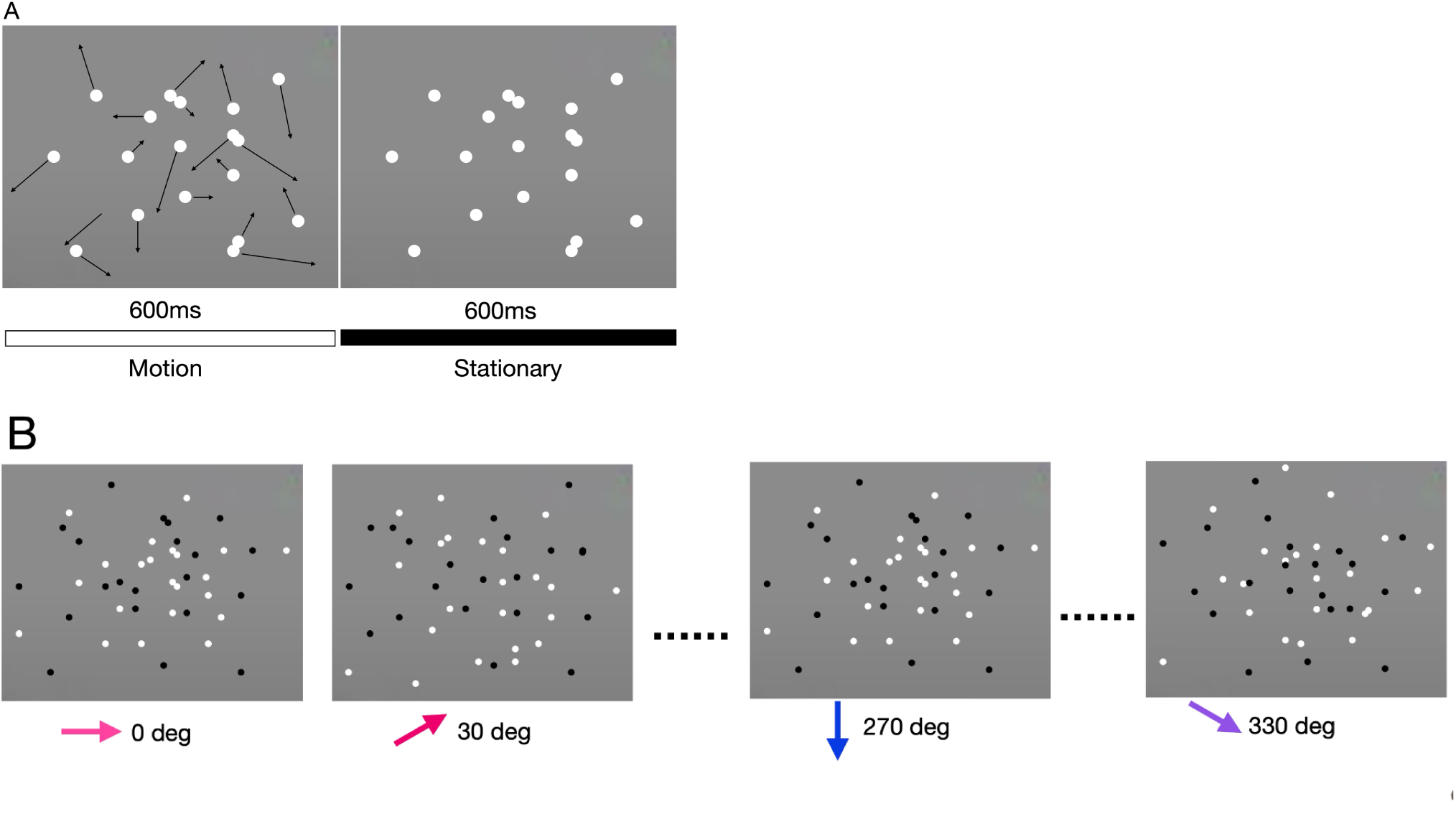
Schematics of MT complex functional mapping stimuli. (A) *Schematics of Moving-stationary dot stimulus for one motion-stationary dot cycle.* One 1.2s long motion-stationary dot cycle consisted of a 600ms period of full-field (incoherent) random dot motion and a 600ms period of stationary dots. The black arrows in the motion-period panel represent each dot’s motion vector over its 50ms lifetime. (B) *Schematics of sequential direction tuning stimulus for one repeat.* In one repeat (24s), coherent full-field dot motion was presented in 12 directions sequentially (0, 30, 60, 90, 120, 150, 180, 210, 240, 270, 300 and 330 degrees) without any inter-stimulus-interval, and each direction was shown for 2s. For both stimuli shown in (A) and (B), the marmosets were freely viewing the motion stimuli without the requirement of fixation.

### Sequential direction tuning stimulus

To investigate if the imaged MT complex revealed the classic physiological properties of (a) a high proportion of individual cells with clear direction selectivity, and (b) functional architecture indicating clustering of direction preference, we presented full-field coherent dot motion (n=300, 150 black and 150 white anti-aliased dots) spanning 12 motion directions (in 30 deg steps) at 16 visual degrees per second (Figure 2B). In each repetition, motion directions were presented sequentially in the counter-clockwise order from 0 degree to 330 degrees, with each direction being presented for 2s. Each block consisted of 20 repeats, with no inter-stimulus-interval/inter-trial-interval between repeats. The positions of all the dots were re-initialized at the first frame of each direction presented. A 4x/0.2NA lens was used for widefield 1P sessions and a 16x/0.8NA lens was used for 2P imaging sessions.

### Data preprocessing

The raw widefield one-photon imaging data were preprocessed with wfield, an open source python package for widefield data analysis and visualization. Specifically, the data collected for widefield microscopy are denoised/decomposed with motion and hemodynamics corrections. The raw two-photon imaging data were preprocessed using Suite2P (Pachitariu et al., 2016, 2018), with a decay constant of 1.8s, to identify regions of interest and extract fluorescence traces for putative cells and their surrounding neuropils.

### Analyses

Analyses for all optical imaging sessions were done in Matlab (Mathworks). For widefield sessions running motion-stationary stimuli and all two-photon imaging data analyses, calcium signals were expressed as dF/F. For each widefield data frame, the dF/F of each pixel within the selected ROI was computed as Formula 1. For each pixel, the ROI’s mean pixel (Froimean) value was subtracted from its raw value (Fpixraw), and then normalized by the ROI’s mean pixel value (Froimean).

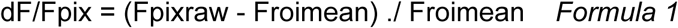

As for two-photon imaging data collected, single cell dF/F was computed as following steps (Formula 2 and 3). First, the dF for each cell (Fcell) was computed by subtracting the average signals of surrounding neuropil with a scaling factor of 0.7 (Keemink et al., 2018; Zong et al., 2022) from its raw signals (Formula 2). Then each cell’s 10th percentile signal was taken as its baseline signal level. Finally, the individual cell’s signal Fcell with the baseline signal F0 being subtracted, was normalized relative to its corresponding baseline (Formula 3).

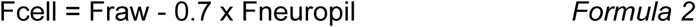

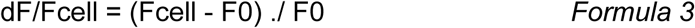

For the widefield data collected for MT complex functional mapping, the data were preprocessed with motion and hemodynamics corrections, and then compressed using singular value decomposition (SVD). Pixelwise activity was reconstructed by multiplying the corrected spatial and temporal components (U × V). Because baseline intensity and slow trends (e.g. hemodynamics) are removed during preprocessing prior to SVD, the reconstructed signals are already zero-centered and represent relative fluorescence fluctuations rather than absolute fluorescence intensity or ΔF/F. All subsequent analyses were performed on the reconstructed signals for widefield direction tuning. The raw data preprocessing and compression were done with wfield, an open source python software package for preprocessing and analyzing widefield (mesoscale) calcium imaging data.

For two-photon imaging data collected for motion direction tuning evaluation, we calculated a direction selectivity index (DSI) for each extracted cell (Formula 4), which is commonly used in electrophysiological studies (Mazurek et al., 2014). A cell’s DSI reflects the difference between its average response signals to the preferred direction (Rpref) and the null direction (Rnull), divided by the sum of its average response signals to the preferred and null directions for normalization. The preferred direction was defined as the motion direction that made the cell elicit the highest average response among the 12 stimulus directions, and the null direction was simply the motion direction that was 180 degrees to the determined preferred motion direction.

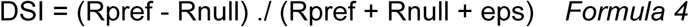

### Brain clearing and lightsheet imaging

After completing the functional calcium imaging data collection, the animals were perfused for histology using 4% PFA administered transcardially. Following perfusion, animals underwent SHIELD fixation, passive delipidation using LifeCanvas delipidation buffer, 2mm sagittal sectioning, and antibodies against GFP and NeuN were delivered using RADIANT stochastic electrotransport primary labeling and secondary stochastic electrotransport labeling to anatomically confirm and quantify the expression (Bürgers et al., 2019; Hillman et al., 2019; Ueda et al., 2020).

### Source, data and code availability

- *Viral Vectors*: The viral vectors were produced by the CPP NIH BRAIN Initiative Armamentarium Vector Core at California State Polytechnic University Pomona: https://www.cpp.edu/armvc/ in coordination with the Beckman Institute CLOVER Center at Caltech https://clover.caltech.edu/.
- *Marmoset brain atlas*: The atlas we used to determine the stereotaxic coordinates of the imaging window center is available online: https://marmosetbrainmapping.org/index.html.
- *Imaging chamber and head restrainment apparatus designs*: The designs for modified imaging chamber, MT-notched halo and headpost are available upon request.
- *Widefield imaging*: The Labcams wfield software packages for widefield imaging acquisition and data preprocessing are available on GitHub repositories: https://github.com/jcouto/labcams, https://github.com/jcouto/wfield. The hemodynamics correction with Labcams integration setup follows the protocol described in ‘Chronic, cortex-wide imaging of specific cell populations during behavior’ (Couto et al., 2021).
- *2-Photon imaging*: 2-Photon imaging data was acquired through Scanbox: https://scanbox.org/, with Neurolabware scope setup: https://neurolabware.com/. The data were preprocessed with Suite2P (https://suite2p.readthedocs.io/en/latest/) (Friedrich et al., 2017; Pachitariu et al., 2016, 2018).
- *Binocular digital DPI eye-tracking*: We acquired eye-tracking data through OpenIris’s DPI plugin. The software is available on GitHub repository: https://github.com/ocular-motor-lab/OpenIris.
- *Brain clearing and lightsheet equipment*: LifeCanvas Technologies: https://lifecanvastech.com/.
- *Experiment stimuli and analysis code with example data:* The Matlab visual stimuli and analysis code with example data is available upon request. All correspondence should be addressed to Dr. Wekselblatt at.

## Results

### Overview

We identified visual area MT (and its satellite regions, together referred to as the MT complex), in both marmosets. Strong and relatively homogenous expression of GCaMP throughout the imaging window allowed us to identify compelling responses to visual motion. This demonstrates the viability of intravenous injections, supporting easier application and more homogenous signals over space across cortical areas. Widefield one-photon imaging revealed strong visual-motion responses and clustered motion direction tuning, with patches of the MT complex showing differential direction preferences. Two-photon imaging further confirmed that the widefield measurements for motion direction tuning are consistent with responses measured at cellular level. Specifically, individual neurons showed tuning for particular directions of visual motion, and were clustered systematically across the imaged area of the MT complex. Together, the functional widefield and two-photon imaging results showed both pervasive direction tuning (a basic hallmark of MT), as well as modular organization of direction tuning. Initially identified by electrophysiological recordings with careful trajectories, the clustering is optical confirmation of the presence of direction columns.

### Assessment of expression and basic physiological identification of visual area MT

To achieve widespread expression of a genetically encoded calcium indicator without local cortical injections, GCaMP8s was packaged into a BBB–crossing AAV capsid (CAP-B10) under the control of the CAG promoter and delivered intravenously to two common marmosets (Figure 1A,B). After an appropriate expression period, a large cranial imaging window was implanted over posterior visual cortex, encompassing the expected anatomical location of area MT (Figure 1C).

Widefield inspection of the cranial window revealed strong and spatially homogeneous fluorescence across the exposed cortical surface, indicating robust expression of GCaMP throughout the imaging field. Fluorescence was stable across imaging sessions and showed minimal spatial heterogeneity, consistent with widespread transduction following intravenous delivery. Imaging through the intact window demonstrated optical clarity suitable for both widefield single-photon imaging and two-photon cellular-resolution imaging, enabling multi-scale functional measurements within the same preparation.

To functionally identify visual-motion–responsive cortex within the imaging window, we performed widefield imaging while the animal viewed full-field dot stimuli that alternated between motion and stationary epochs (Figure 2A). These stimuli elicited robust fluorescence responses across posterior extrastriate cortex. Among several visually responsive regions, one contiguous region (ROI1) exhibited consistently stronger and more reliable modulation by visual motion. Within this region, motion-evoked responses were spatially coherent across smaller subregions and exhibited elevated activity during motion epochs and reduced activity during stationary epochs (Figure 3B–D). This response profile is consistent with classic electrophysiological descriptions of MT neurons, which show elevated firing rates during visual motion and reduced activity in the absence of motion. Based on its anatomical location and functional response properties, this region was identified as the MT complex (K. H. Britten et al., 1993; Gharaei et al., 2013; Lui & Rosa, 2015; Smith et al., 2005; Snowden et al., 1992; Tsui et al., 2010).

**Figure 3.**
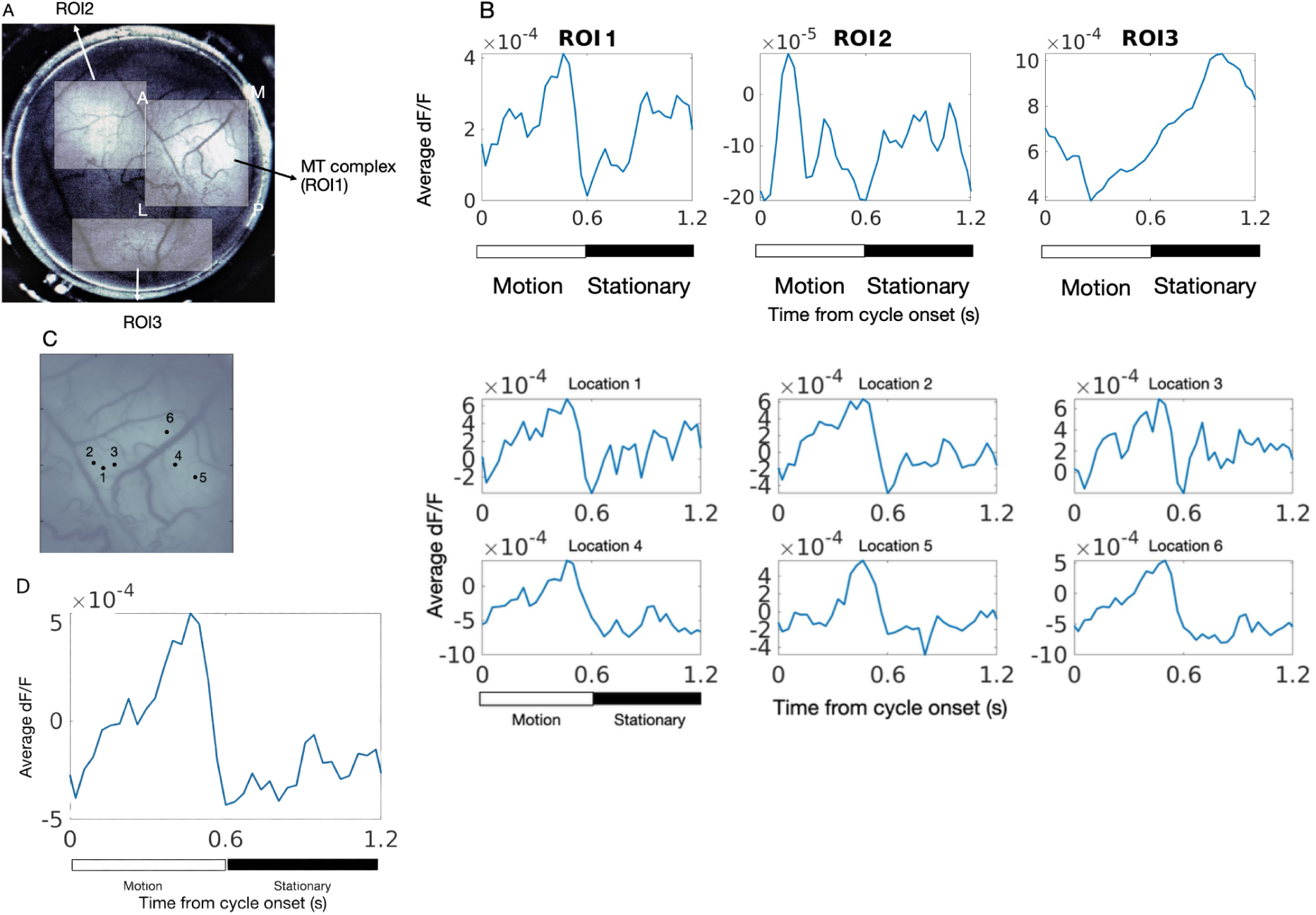
Functional identification of MT complex following systemic GCaMP expression with motion-stationary dot stimulus. (A) *Visually responsive ROIs identified under 2x lens*. Through 40 trials of stimuli that presented 800 cycles of motion-stationary dots, on average, there were three regions consistently showed visually evoked responses: ROI1: 3.77mm x 3.05mm, ROI2: 3.23mm x 2.88mm, ROI3: 1.80mm x 3.59mm. (B) *Average dF/F traces of the three ROIs over 800 motion-stationary cycles*. (C) *Average activity of 6 example locations selected within MT complex (ROI1) to 800 motion-stationary dots cycles.* Left: Selected locations overlaid with the mean response image of MT complex/ROI1. Each location is 10×10 pixels in size, approximately 0.18×0.18mm. Right: The average dF/F traces of the six selected locations. (D) *Average dF/F traces over the 6 example locations selected within the MT complex*.

### Widefield and two-photon measurements reveal canonical direction tuning in MT

Having identified the MT complex, we next characterized motion direction tuning at both population and cellular levels using full-field coherent dot motion presented sequentially in 12 directions (Figure 2B). During widefield imaging sessions, approximately a 4.75 × 4.75 mm cortical area was imaged under a 4× objective.

Motion-evoked GCaMP signals were corrected for motion artifacts and hemodynamic contributions prior to analysis. Average response images revealed a region of strong stimulus-driven GCaMP activity that overlapped closely with the MT complex identified using motion–stationary stimuli (Figure 4A). In contrast, hemodynamic signals exhibited broader, more spatially uniform responses.

**Figure 4.**
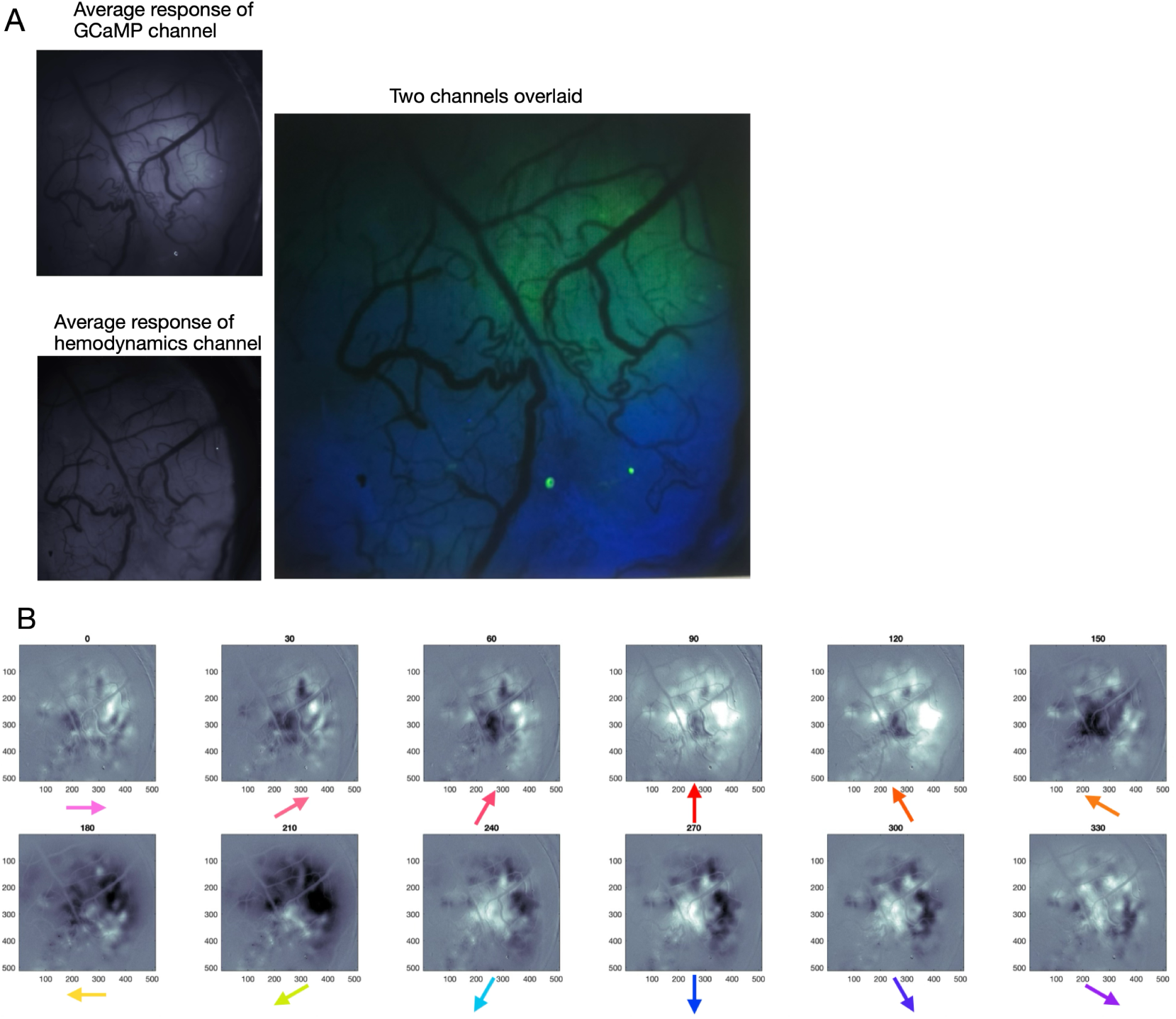
Direction preference patches in marmoset MT complex revealed by widefield one-photon imaging. (A) *Functional GCaMP responses vs Hemodynamics responses.* Left: top panel shows the average response image of the functional GCaMP channel, and the bottom panel shows the average hemodynamics response image of the hemodynamics correction channel. Right: Two channels average responses are overlaid. Green indicates the specific area was functionally responsive to visual-motion stimuli and the responses were mainly driven by GCaMP. Whereas blue suggests the activity/expression was more driven by slow hemodynamics signals. The 4x lens covered approximately 4.57×4.57mm large cortical area. (B) *Average response images of the MT complex for 12 motion directions.* Each row covers 180 degrees of motion direction, and each column shows a pair of directions that are 180 degrees opposite from each other.

Population-level direction tuning maps demonstrated that different patches of cortex within the MT complex exhibited distinct preferred motion directions (Figure 4B). Direction preference varied smoothly across the cortical surface, forming spatially organized domains. Motion directions separated by 180° preferentially activated largely nonoverlapping cortical regions, and direction sets spanning opposite halves of the stimulus space (0–180° vs. 180–360°) engaged distinct cortical slabs. These widefield measurements provide a continuous spatial view of direction preference organization that would be difficult to obtain using sparse electrode sampling and are consistent with the existence of direction columns in primate MT inferred from electrophysiological studies.

### Cellular confirmation of direction tuning with two-photon imaging

To validate population-level measurements at cellular resolution, we performed two-photon imaging in multiple regions of interest within the MT complex using the same direction-tuning stimuli (Figure 5). Each two-photon session imaged an approximately 800 × 800 µm field of view using a 16× objective.

**Figure 5.**
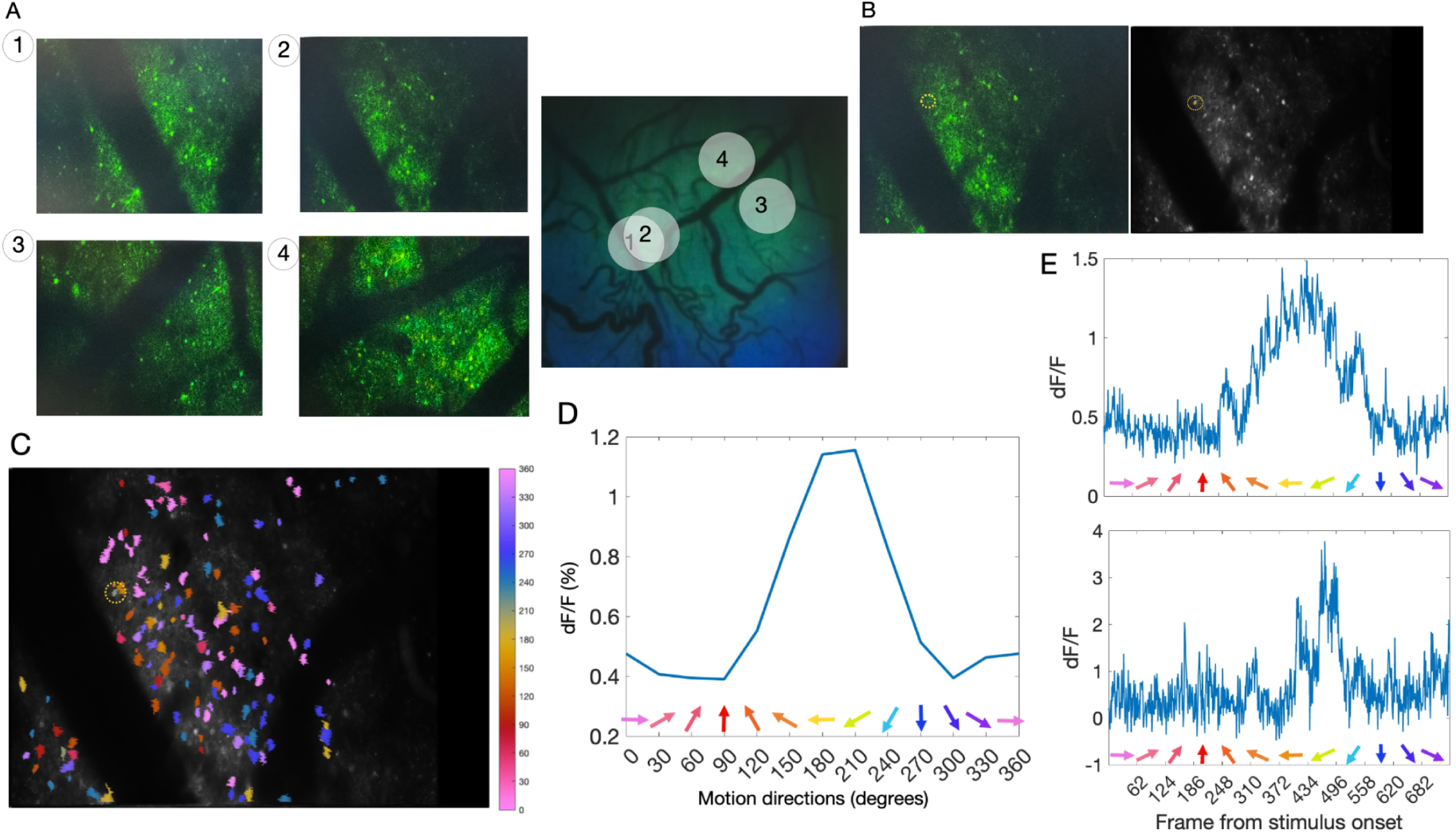
Single cell direction tuning in marmoset area MT measured by two-photon imaging. (A) *Example 2P sessions and their imaging locations.* Left: four 2P sessions’ example recording frame snapshots with green labeling GCaMP expression. Right: four 2P sessions’ imaged areas (shown on the left) overlaid with 1P image from Figure 4A-right panel. The example 2P session of area 2 is the focus of the following analyses in the report. The session of area 4 is the last imaging session from Animal1, 13 months after tail vein injection. (B) *2P area 2’s session recording frame image (Left) and the session mean image (right).* The example cell’s location is outlined by the dashed-circle, and the cell’s mask is colored in the mean image. (C) *Single cell direction tuning.* The cells with direction selectivity index higher than 0.05 (DSI>0.05, n=138) are filled and labeled with colors indicating their motion direction preferences. The colorbar’s color coding for each direction is consistent with that of the direction arrows in the other plots. The example cell is outlined with a dashed-circle. (D) *Example cell’s direction tuning.* The line plot shows the direction tuning curve of the example cell highlighted in (B) and (C). (E) *Example cell’s responses to 12 motion directions.* Top: The calcium signal (dF/F) as a function of frames averaged across 20 repeats. Bottom: The calcium signal (dF/F) of an example single run through motion directions (in this case, repetition #14).

Across four example imaging locations distributed within the MT complex, two-photon images revealed robust GCaMP expression in individual neurons and surrounding neuropil (Figure 5A). The locations of these two-photon recordings were overlaid onto widefield functional maps, enabling direct comparison between population-level and cellular-level measurements. Subsequent analyses focused on one representative imaging region (area #2), in which 170 neurons exhibited stimulus-driven responses.

Direction selectivity indices (DSIs) were computed for all responsive neurons, and neurons with DSI > 0.05 (n = 138) were classified as direction selective. These neurons exhibited a range of preferred motion directions that reflected the local direction preference observed in widefield maps (Figure 5C). Clusters of neurons sharing similar direction preferences were observed, with gradual transitions in preferred direction across the imaging field, consistent with columnar organization.

Example direction tuning curves from individual neurons demonstrated broad, stereotypical MT-like tuning, with peak responses at the preferred direction and reduced responses at nonpreferred directions (Figure 5D,E). These tuning properties were evident both in trial-averaged responses and in individual stimulus repetitions, indicating robust single-neuron direction selectivity with moderate trial-to-trial variability.

### Longitudinal stability of functional imaging signals

Across 13 two-photon imaging sessions conducted in one animal’s MT complex, we identified a total of 1,336 visually responsive neurons, with approximately 100 neurons per session. Direction tuning properties were consistent across sessions and aligned with canonical MT physiology (Albright, Desimone and Gross, 1984).

The final two-photon imaging session, conducted 13 months after viral injection and immediately prior to brain clearing and light-sheet imaging, revealed robust motion-driven responses and stable GCaMP expression comparable to earlier sessions. These observations indicate that systemic AAV delivery can support long-term, longitudinal functional imaging in the marmoset cortex.

### Histological confirmation of expression using tissue clearing and light sheet microscopy

Following completion of in vivo imaging experiments, we performed ex vivo tissue clearing and light-sheet microscopy to assess the anatomical distribution of systemically delivered GCaMP expression across the brain (Figure 6). Cleared hemispheres were imaged volumetrically to visualize expression throughout cortex and subcortical structures (Keller & Ahrens, 2015; Qu et al., 2022, 2022; F. Xu et al., 2021). The speed of light sheet imaging and much lower hands-on time for tissue processing are important factors to consider in larger animal species, and development of better clearing and staining techniques in large tissues is of great interest to us.

**Figure 6.**
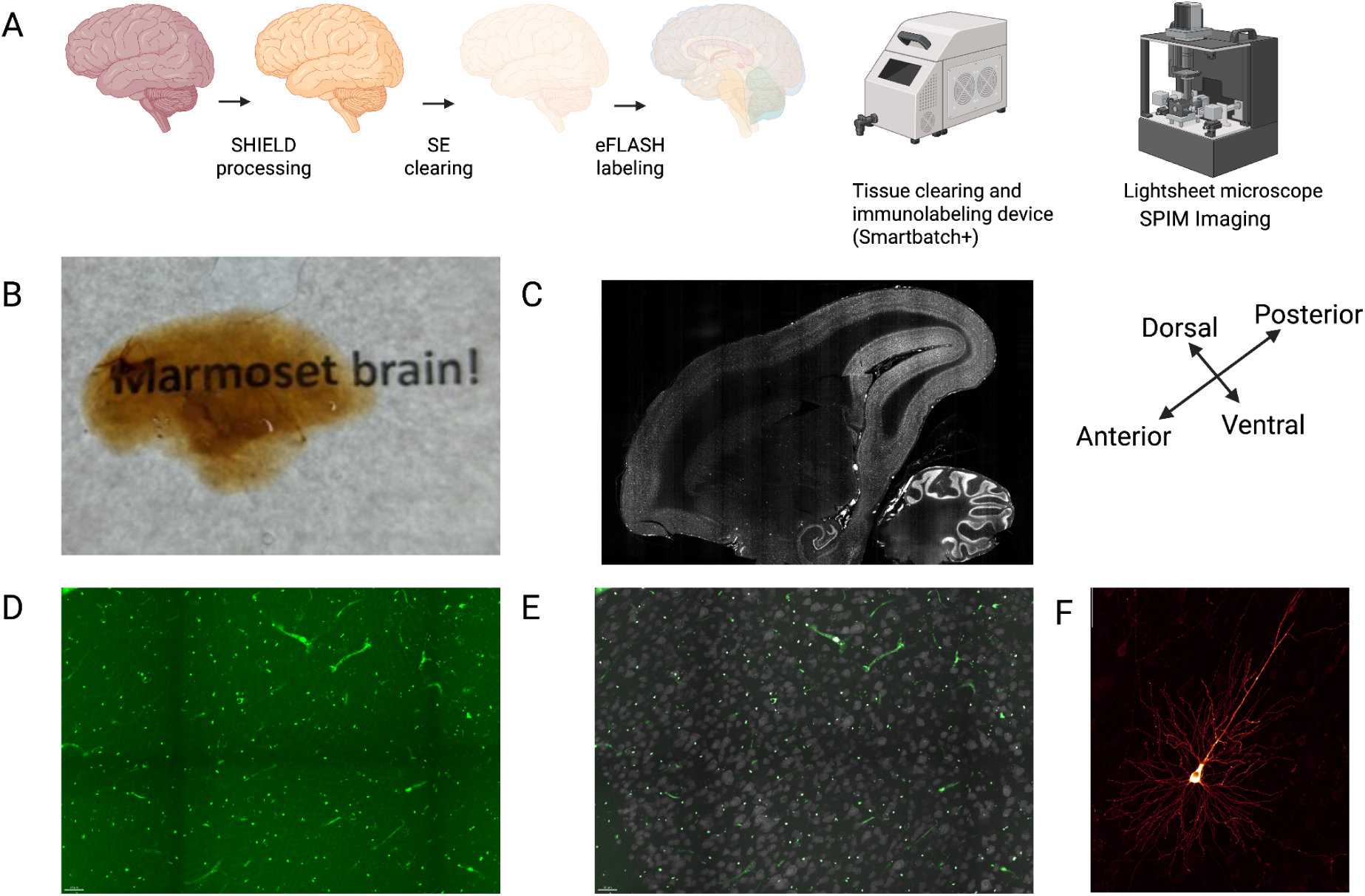
Assessment of expression using tissue clearing and light sheet microscopy. (A) Schematic of brain clearing and staining procedure using LifeCanvas Technologies system. Cleared Tissue is immunostained and then imaged using a lightsheet microscope. High resolution imaging is performed on Andor spinning disk confocal. (B) Whole marmoset hemisphere after clearing procedure. Text is visible through the cleared sample. (C) 2mm thick sagittal section of marmoset cleared brain showing NeuN staining of neuronal cell bodies - imaged with 3.6x objective on the light sheet microscope. (D) 30x objective on spinning disk confocal of visual cortex neurons stained with anti-GFP showing GCaMP expression (green) in cleared tissue. 20um max intensity projection (MIP) (.2um, .2um, 1um) (x,y,z). (E) 30x of the same visual cortex region as (D) but showing the NeuN label of cell bodies in white, as in (C). (F) 30x zoom in on a single neuron stained with V5 antibody in the marmoset cortex expressing smGFP-V5 after delivery of CAP-B10-CAG-smV5-GFP. (construct shown in Supplemental Figure 4).

Hemispheric reconstructions revealed widespread GCaMP expression across cortical areas, consistent with the uniform fluorescence observed through the cranial window during in vivo imaging and with prior reports of CAP-B10-mediated systemic delivery. Expression was broadly distributed across the cortical sheet rather than localized to discrete injection sites, confirming that intravenous administration resulted in large-scale neuronal transduction.

Because native GCaMP fluorescence provided limited anatomical contrast in cleared tissue, we additionally evaluated a companion CAP-B10 construct expressing a membrane-localized, epitope-tagged GFP reporter to enhance visualization of neuronal morphology (Hillman et al., 2019; Ueda et al., 2020). Immunolabeling of this reporter yielded higher signal-to-noise and clearer anatomical labeling while exhibiting a spatial distribution consistent with that observed for GCaMP expression (Figure 6F). Together, these measurements provide anatomical confirmation that systemic CAP-B10 delivery produces widespread cortical labeling suitable for functional imaging.

Both the mouse and marmoset preparations revealed the GCaMP8 native fluorescence is lower signal than previous versions of GCaMP(6-7), as well as compared to eGFP, preventing high signal-to-noise measurements of anatomical features of neurons such as synaptic connections. For this reason we also used a similar construct with the CAP-B10 capsid carrying the same promoter (CAG), but this time driving a membrane localizing V5-tagged spaghetti monster green fluorescent protein (smV5-GFP) with a CAAX motif on the C-terminal end and 10 V5 tags for enhanced anatomical reconstructions using light sheet microscopy as seen in figure 6F (Akerboom et al., 2012; Tian et al., 2009, 2012; Zhang & Looger, 2024). We found that the epitope tags on the spaghetti monster construct allow much finer immunolabeling in the cleared tissue and yielded much higher signal in expressing neurons for downstream analyses - confirming the expression patterns seen with the GCaMP carrying CAP-B10.

## Discussion

Across two marmosets, we combined widefield single-photon imaging, two-photon cellular-resolution imaging, histological sectioning, and whole-brain clearing with light-sheet microscopy to validate systemic AAV-mediated delivery of a genetically encoded calcium indicator (GCaMP8s). Functional measurements were performed in and around the primate visual area MT, allowing us to validate the measured signals by leveraging well-established physiological properties of this area, such as strong responses to visual motion and robust direction tuning at both population and cellular levels. Together, these results indicate that intravenous delivery supports stable and relatively homogeneous expression suitable for quantitative functional imaging across spatial scales in the primate cortex.

To our knowledge, relatively few studies have provided functional validation of GCaMP delivered via blood–brain barrier–crossing AAVs in the cortex of awake, behaving non-human primates. In this study, we show that (1) the CAP-B10 capsid supports systemic neuronal labeling in the marmoset brain, (2) systemic delivery of GCaMP via CAP-B10, in combination with a chronic imaging chamber, enables longitudinal functional imaging, and (3) activity measured using this approach yields estimates of stimulus-driven responses and functional organization that are commensurate with those obtained using electrophysiological methods, at least in the primate visual area examined here (the MT complex).

### Comparison with electrophysiological measurements of MT

The primate MT complex has been studied extensively using single-unit and multi-unit electrophysiology, yielding detailed characterizations of motion direction selectivity, speed tuning, spatiotemporal receptive field structure, and columnar organization (Born & Bradley, 2005; Huang et al., 2025; Kumano & Uka, 2023; J. Liu & Newsome, 2003; Lui & Rosa, 2015). This body of work has established MT as a solid reference point for evaluating new measurement approaches. Consistent with this literature, our widefield single-photon and two-photon imaging measurements revealed strong motion-driven responses, robust direction tuning at the single-neuron level, and clustered spatial organization of direction preferences that varied systematically across the cortical surface.

While electrophysiological recordings provide high temporal resolution and direct measurements of spiking activity, they necessarily sample a limited subset of neurons and offer an incomplete view of functional organization across large cortical regions (Cybulski et al., 2015; Kaszás et al., 2025; Levina et al., 2022). Optical imaging approaches, by contrast, enable simultaneous measurements across large neuronal populations and millimeter-scale cortical areas. The close correspondence between imaging-based measurements reported here and established electrophysiological descriptions of MT supports the use of systemically delivered calcium indicators for studying functional organization in the primate cortex, while underscoring the complementary strengths of these modalities.

### Advantages of systemic over local viral injection strategies

Most prior optical imaging studies in nonhuman primates have relied on local intracortical viral injections to express genetically encoded indicators (Campos et al., 2023b; Heider et al., 2010; Seidemann et al., 2016). Although effective, this approach introduces several constraints, including the need for multiple cortical penetrations, spatially heterogeneous expression, and variability across injection sites (Bohlen & Tremblay, 2023; Federer et al., 2024; Kojima et al., 2021). These factors complicate widefield imaging and can limit the interpretability of large-scale population measurements (Matsui et al., 2024; Nietz et al., 2022b; O’Shea et al., 2017).

In contrast, systemic intravenous delivery using a BBB-crossing capsid produced robust and relatively uniform expression across the cortical surface without the need for local cortical injections. This homogeneity is particularly advantageous for widefield imaging, where continuous spatial patterns of activity are of interest. In addition, avoiding cortical injections simplifies surgical preparation and facilitates longitudinal imaging across extended time periods by avoiding mechanical damage and immunogenicity in the imaging zone, making this approach well suited for repeated measurements and large-scale functional mapping.

### Multi-scale validation of functional signals

A key feature of the present study is the combined use of widefield single-photon imaging and two-photon cellular-resolution imaging within the same animals. Widefield imaging provided a continuous view of functional organization across the MT complex, revealing large-scale direction tuning patterns consistent with columnar structure. Two-photon imaging within selected regions of interest further demonstrated that these population-level patterns reflected the tuning properties of individual neurons. The concordance between widefield and cellular measurements supports the interpretation of widefield signals in terms of underlying neuronal activity.

### Limitations and future directions

Several limitations should be considered when interpreting these results. While the toolkit of cell type specific enhancers for NHPs continues to improve and expand, systemic delivery does not offer the same degree of spatial or cell-type specificity as targeted local viral injections (Bourdenx et al., 2014; Deng, 2010; Serwer et al., 2010; Zhou et al., 2022), which may be preferable for experiments requiring precise genetic or anatomical control. That being said, newly developed enhancer AAVs show remarkable cell-type specificity when injected in rodents and NHPs. Direct injection of high doses of enhancer AAVs leads to a loss of cell-type specificity, favoring systemic injection approaches where there is a lower multiplicity of infection and regulatory design features behave with high fidelity (Hunker et al., 2024; Kussick et al., 2024). In addition, calcium imaging provides an indirect measure of neuronal activity and lacks the temporal precision of electrophysiological recordings (Chung et al., 2019; Obien et al., 2015; Paulk et al., 2022; Trautmann et al., 2023).

Future studies combining optical imaging with multi-unit or laminar electrophysiological recordings within the same cortical regions may provide a more complete characterization of functional organization. Finally, the present study was conducted in a limited number of animals, reflecting the technical demands of primate imaging, and future work will be important for assessing variability across individuals and preparations.

Additionally one may consider combining functional indicators with high-contrast anatomical reporters may facilitate integrated in vivo imaging and ex vivo reconstruction within the same preparation. For example, packaging both GCaMP and smV5-GFP in the same capsid to allow for functional imaging in-vivo, as well as accurate and high resolution reconstructions of the same cells ex-vivo in full 3D volumes of cleared tissue.

In summary, these results indicate that systemic AAV delivery can support stable, multi-scale functional imaging in the marmoset brain. By demonstrating close correspondence between imaging-based measurements and established electrophysiological descriptions of the MT complex, this work suggests that intravenous delivery of genetically encoded calcium indicators provides a practical complement to existing electrophysiological and local viral approaches for studying functional organization in the primate cortex. The present study focuses on validating the feasibility and fidelity of systemic AAV-mediated functional imaging; more detailed analyses of MT functional architecture are reserved for a subsequent report.

## Acknowledgements

This publication was supported by and coordinated through the BRAIN Initiative Armamentarium for Precision Brain Cell Access. We thank Dr. João Couto for setting up Labcams software for 1P-widefield imaging acquisition and data preprocessing, Dr. Dario Ringach and Dr. Joshua Trachtenberg for setting up Neurolabware equipment and Scanbox for 2-Photon imaging data acquisition, Dr. Jude Mitchell for helping us design the stimuli for MT functional mapping, Hsuan Lee and Zachary Woods from LifeCanvas Technologies for brain clearing and lightsheet imaging development, and Jose Gomez Godinez for fabricating imaging chamber apparatus and head restraint parts.

Marmoset work was done in collaboration with the Fuster Behavioral Operations and Innovation Core at UCLA, providing helpful advice and experimental expertise for marmoset work, and pUCmini-iCAP-AAV.CAP-B10 was a gift from Viviana Gradinaru (Addgene plasmid # 175004; http://n2t.net/addgene:175004; RRID:Addgene_175004)

