## supplemental figures for "Systemic AAV delivery of a calcium indicator in marmosets: functional validation in visual area MT"

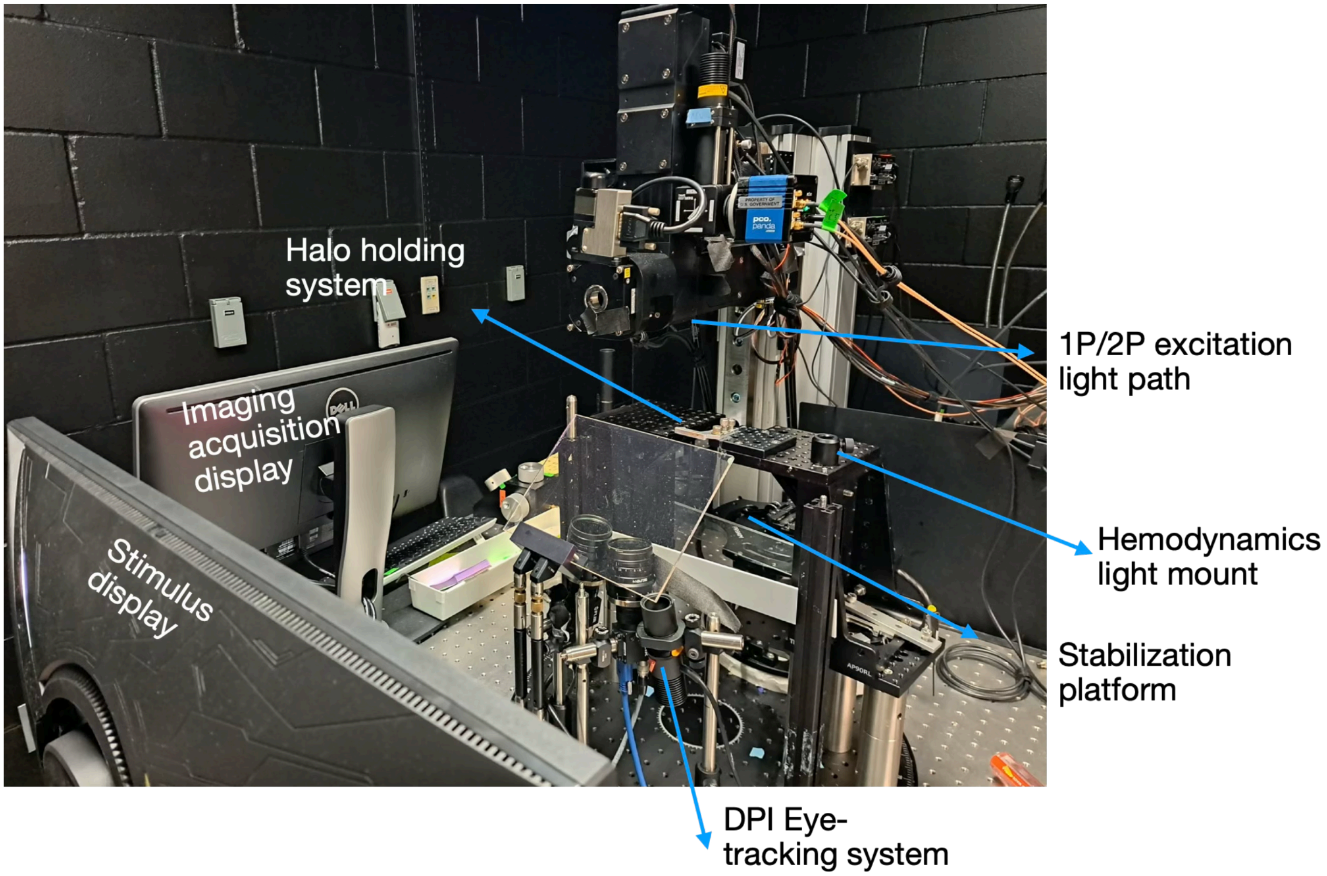


**Supplemental Figure 1:** Experiment platform for widefield and two-photon imaging in awake marmosets.

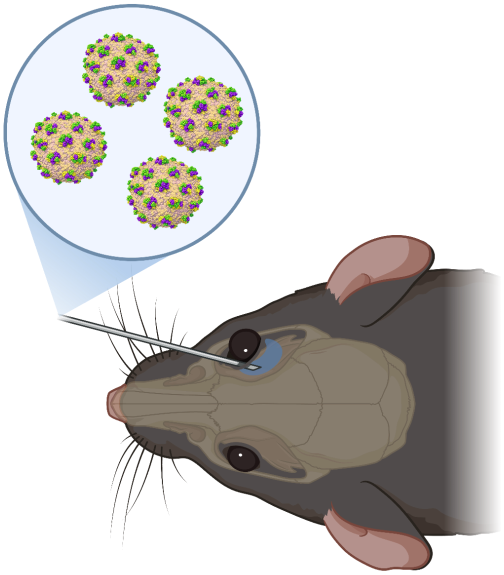

**Supplemental Figure 2:** Schematic of retro-orbital injection in mice.

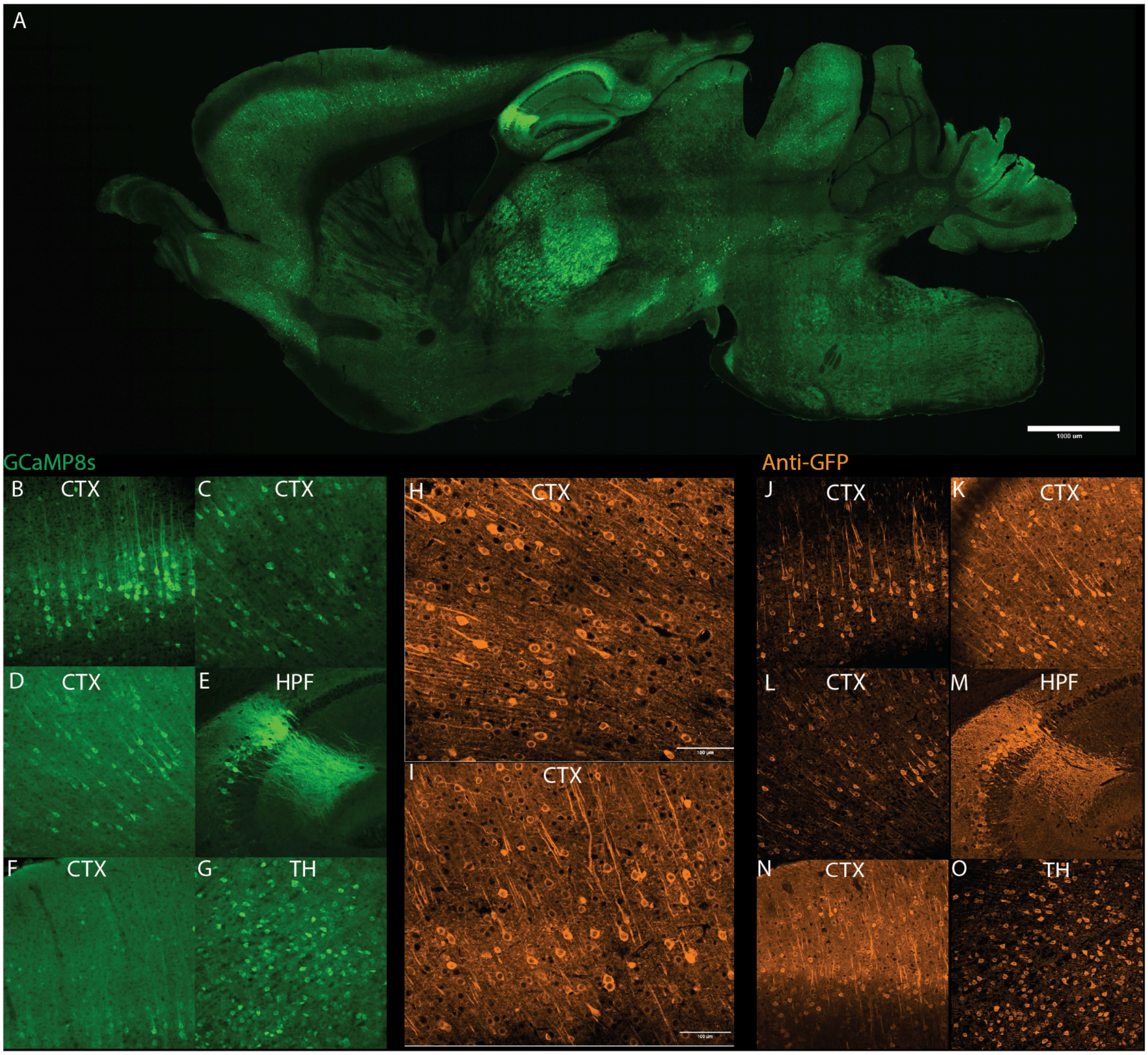


**Supplemental Figure 3:** Histology of a sagittal section (100um thickness) from a C57BL/6J mouse injected with 3E11 vg retro-orbitally with CAP-B10-CAG-GCaMP8s following two weeks of incubation. (A) Low magnification image (20x objective) of endogenous green fluorescence from GCaMP8s in a sagittal section (scale bar is 1000um). (B-G) Green fluorescence signal in cortex (B-D), Hippocampus (E), cortex (F), and thalamus (G). GFP-antibody labeling at high magnification (40x objective) (H) and (I) in cortex (scale bar, 100um). 20x images of GFP antibody labeling in cortex (J, K, L), hippocampus (M), cortex (N), and thalamus (O)

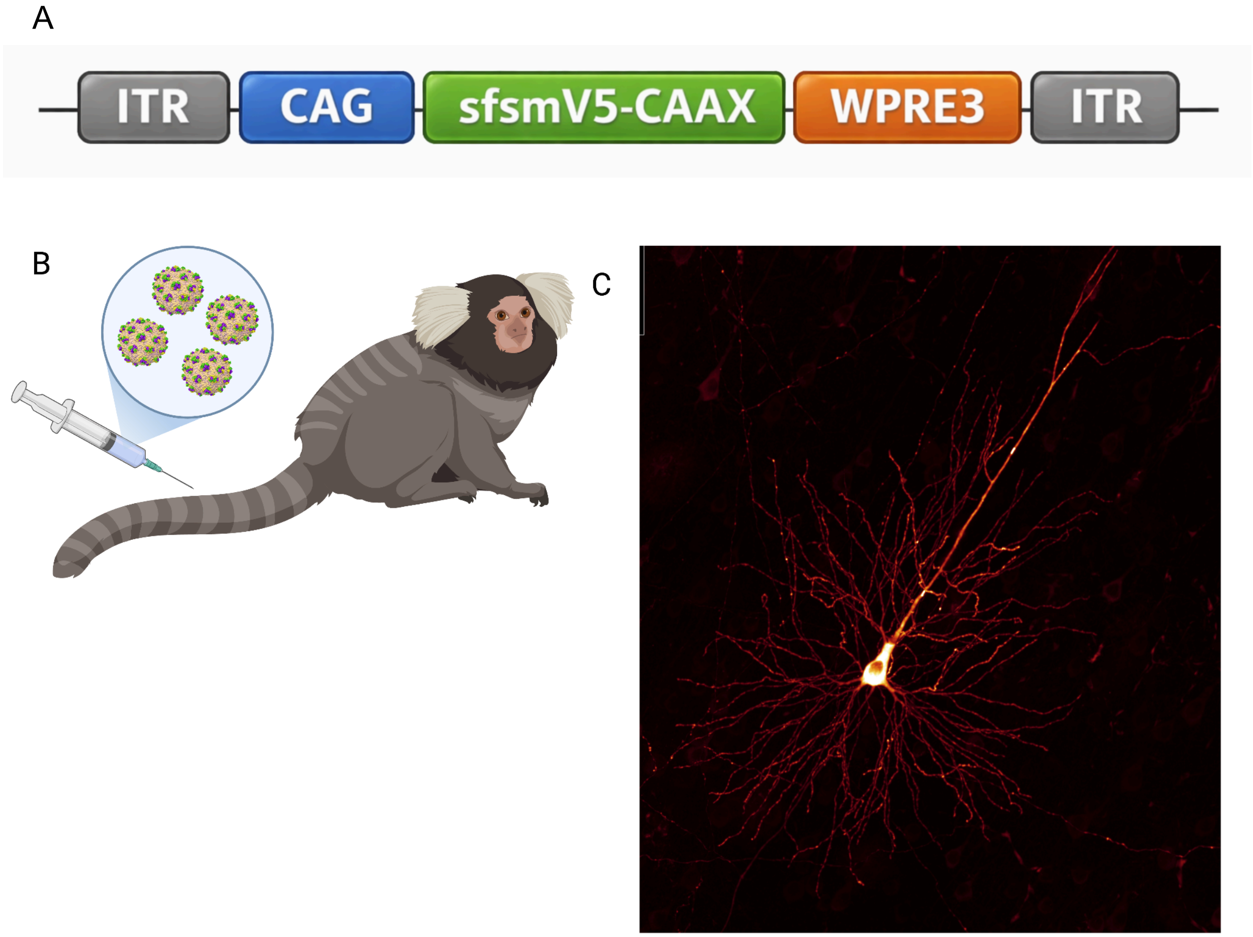


**Supplemental Figure 4:**
(A) Schematic of the viral construct used in this study (AAV9-CAP-B10-CAG-smV5-GFP-CAAX ).
(B) Schematic of intravenous delivery of the BBB-crossing capsid via tail-vein injection.
(C) 30x zoom in on a single neuron in the marmoset cortex expressing smGFP
